# The African Bullfrog (*Pyxicephalus adspersus*) genome unites the two ancestral ingredients for making vertebrate sex chromosomes

**DOI:** 10.1101/329847

**Authors:** Robert D. Denton, Randal S. Kudra, Jacob W. Malcom, Louis Du Preez, John H. Malone

## Abstract

Heteromorphic sex chromosomes have evolved repeatedly among vertebrate lineages despite largely deleterious reductions in gene dose. Understanding how this gene dose problem is overcome is hampered by the lack of genomic information at the base of tetrapods and comparisons across the evolutionary history of vertebrates. To address this problem, we produced a chromosome-level genome assembly for the African Bullfrog (*Pyxicephalus adspersus*)—an amphibian with heteromorphic ZW sex chromosomes—and discovered that the Bullfrog Z is surprisingly homologous to substantial portions of the human X. Using this new reference genome, we identified ancestral synteny among the sex chromosomes of major vertebrate lineages, showing that non-mammalian sex chromosomes are strongly associated with a single vertebrate ancestral chromosome, while mammals are associated with another that displays increased haploinsufficiency. The sex chromosomes of the African Bullfrog however, share genomic blocks with both humans and non-mammalian vertebrates, connecting the two ancestral chromosome sequences that repeatedly characterize vertebrate sex chromosomes. Our results highlight the consistency of sex-linked sequences despite sex determination system lability and reveal the repeated use of two major genomic sequence blocks during vertebrate sex chromosome evolution.

## MAIN TEXT

Sexual reproduction is the dominant mode of creating offspring among vertebrates^1^, but the fundamental genetic mechanisms that produce two sexes are—in contrast—staggeringly diverse^2,3^. This diversity of genetic sex determination systems is what dictates the formation of sex chromosomes, which differ widely from autosomal chromosomes in their lability, gene content, and structural organization^4,5^. A particular departure from autosomal characteristics is the phenomenon of heteromorphic sex chromosomes which differ in size and gene content. As sex chromosomes evolve, the sex chromosome associated with the heterogametic sex (either Y or W) can experience gene loss, rearrangements, and gains of heterochromatin that change chromosome size and reduce gene dosage on the gametologous X or Z chromosome, respectively^6–12^.

At the same time, reductions in gene dosage are largely deleterious^13–17^. Mechanisms of dosage compensation, such as X-inactivation in eutherian mammals^18–20^ and X-linked hyperexpression in lizards^21,22^ are thought to help solve this problem. However, sex-linked genes are expressed at levels proportional to their copy number in birds, snakes, and liver flukes—all species with ZW sex determination—suggesting a lack of dosage compensation^6,23–27^. The ability to compensate gene dose reductions during sex chromosome evolution in some lineages but tolerate gene dose reductions in others remains poorly understood^24,28^. One hypothesis that addresses this phenomenon is that certain genomic regions are more dose tolerant than others, and shuffling of these syntenic blocks has contributed to the repeated evolution of heteromorphic sex chromosomes^28–30^.

The diversity of vertebrate sex determination systems provides a significant obstacle for understanding the gene dose challenges created during sex chromosome evolution. Mammals and birds each have a single sex determination system (XY and ZW, respectively), whereas multiple sex determination systems have evolved independently in fish, amphibians, and non-avian reptiles^2^. The majority of fish, amphibians, and reptiles with genetic sex determination have homomorphic sex chromosomes, supporting the idea that repeated sex chromosome turnover eliminates gene dosage imbalance^31^. However, a key piece of missing information is the content and evolutionary origin of heteromorphic sex chromosomes at the base of Tetrapoda, which could identify the frequency that ancestral chromosome segments have become sex-linked and dosage imbalanced and permit comparisons across the evolutionary history of vertebrates. Here, we provide the first chromosome-level genome assembly of an amphibian with heteromorphic sex chromosomes (the African Bullfrog, *Pyxicephalus adspersus*). We identified and validated both the Z and W sex chromosomes and compared sequences with reduced gene dosage across the evolutionary history of vertebrates. By filling in this major gap in amphibians, we find that sequences that evolve to be on heteromorphic sex chromosomes can be primarily traced back to two ancestral vertebrate chromosomes, one of which is more dosage sensitive than the other, supporting the hypothesis that there are limited options for evolving vertebrate heteromorphic sex chromosomes^28,29^.

## RESULTS

### Genome Assembly

To identify sex-linked sequences in the bullfrog, we extracted DNA from both a female and a male individual from inbred captive lines and sequenced the genome to 138X coverage using Illumina HiSeq 2000 paired-end reads (Supplementary Table S1). We then assembled reads from the female into a preliminary genome assembly using ALLPATHS-LG^32–34^. This assembly spanned ~1.56 GB, consistent with previous estimates of genome size for this species^35,36^. Next, Chicago and HiC long-range libraries were prepared and sequenced by Dovetail Genomics (Santa Cruz, CA) and combined with our draft assembly and high coverage fragment libraries to provide an addition 51X coverage and generate a final assembly using the HiRise scaffolder^37^.

The genome was assembled into 5,411 scaffolds, had an N50 of 157.52 Mb, and contained 91.7% of the benchmarking single copy orthologs from eukaryotes^38^ (Supplementary Table S1). Fourteen of the assembled scaffolds were exceptionally large compared to all others and contained 99% of the total assembly. The mean scaffold length of these 14 scaffolds was 110 Mb, an order of magnitude larger than the 4.2 kb average size for the remaining 5,397 scaffolds. The dramatic difference between the size of the largest scaffolds compared to all others suggested the possibility that these scaffolds represented fully assembled chromosomes. We compared the sizes of the 14 scaffolds to the chromosome number and sizes observed from African Bullfrog karyotypes. The chromosome number observed from karyotype analysis of sequenced individuals was consistent with prior observations^35,36^: 12 autosomes plus a heteromorphic Z/W, for a total of 14 chromosomes (Figure 1A). The DNA sequence sizes of the 14 scaffolds matched very closely to the lengths of chromosomes as measured from karyotypes (*R*^2^ =0.93; Figure 1B). Taken together, the exceptional size, totality of sequence, and high similarity to karyotype number and lengths support that these 14 scaffolds are assembled chromosomes—the first for a heteromorphic amphibian, and only the third amphibian assembled at chromosome-scale^39,40^.

**Figure 1.**
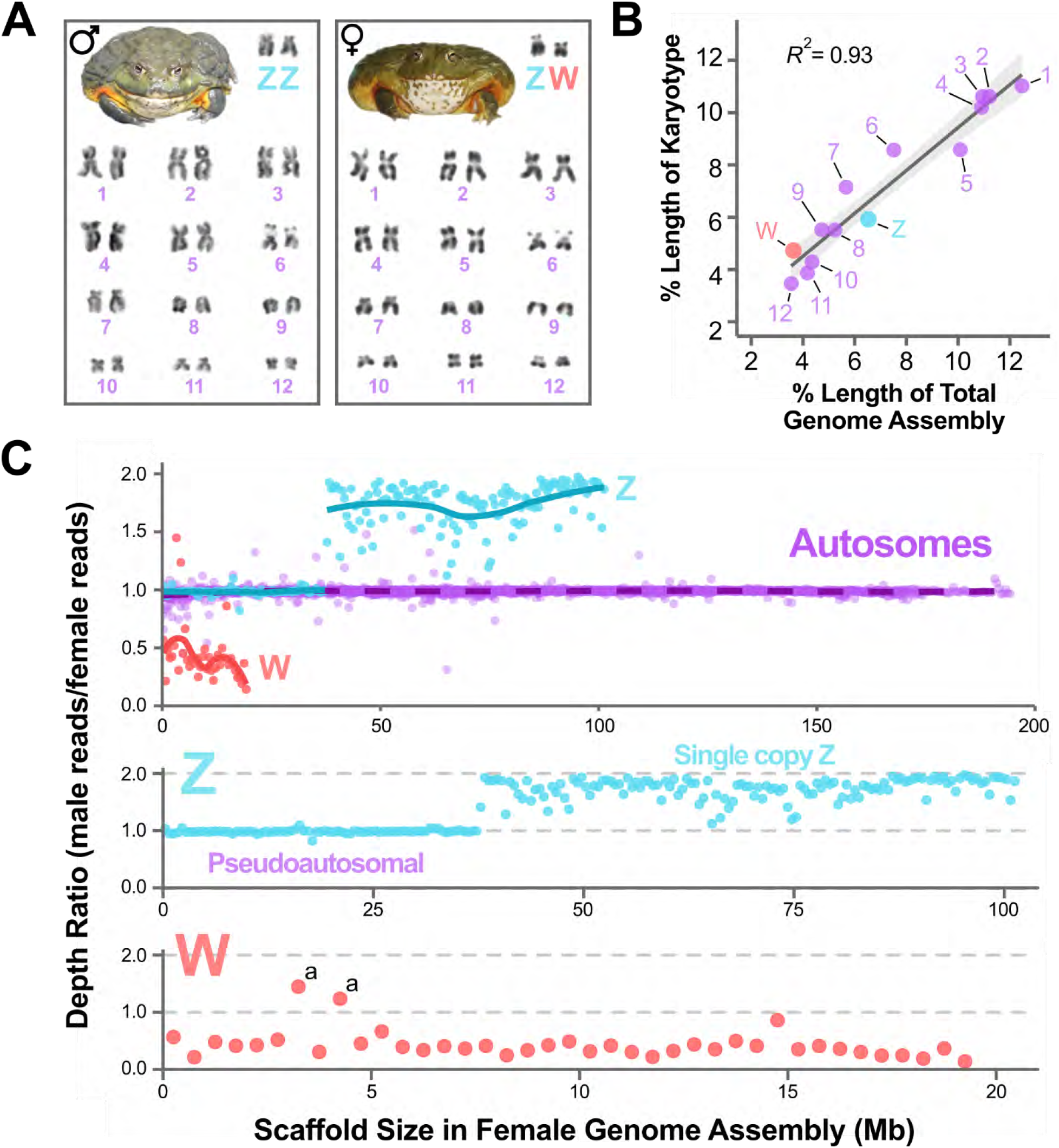
The genome of the African Bullfrog (Pyxicephalus adspersus). **(A)** Females are the heterogametic sex, as observed by the heteromorphy between the Z and W chromosome from karyotype analysis and consistent with previous observations^35,36^. **(B)** Fourteen scaffolds contain ~99% of the African Bullfrog genome assembly, and these scaffolds matched both the expected number (N = 14) and size distribution of chromosomes based on karyotypic measurements. **(C)** These 14 chromosome scaffold were assigned as autosomal (purple), Z (blue), or W (red) based on the proportion of male and female reads that were mapped onto the female genome assembly (total number of reads = 4 x 10^9^). Each dot represents the depth ratio (male/female) averaged across a 500 kb sliding window, and the male depth is multiplied by 0.97 to account for the greater number of raw reads produced for the male individual (top panel). Solid lines are smoothed conditional means created using a general additive model (GAM) function in R^41^. Because African Bullfrogs have heteromorphic sex chromosomes, depth ratios for sex chromosomes were expected to be near 2 (Z-linked) or 0 (W-linked). Shared Z-W sequences and autosomal scaffolds were expected to display depth ratios of 1. The bottom two panels display the same 500 kb windows for the identified Z and W, respectively. Outlier windows along the W scaffold (marked with asterisks) include genes that are likely inflated due to the presence of genes with multiple copies located on autosomes (a: 28S, 18S).

### Identification of Sex-linked Scaffolds

To assign each of the large scaffolds as either autosomal or sex chromosomes, we mapped all the reads from both the male and female back to the female assembly (Supplementary Table S2). Because sex chromosomes have different copy numbers in African Bullfrogs, we expected autosomal scaffolds to have equal coverage between the sexes, Z-linked scaffolds to have 2-fold higher coverage in the male, and W-linked scaffolds to have coverage for females and very low or zero coverage for males. Using this approach, we identified 1 Z, 1 W, and 12 autosomal scaffolds (Figure 1C). The single Z-assigned scaffold, which measured 92.5 Mb in size, accounted for 91% of the predicted size of bullfrog Z as measured from karyotypes. The Z-assigned scaffold included a ~37 Mb region that displayed the same 1:1 coverage ratio as autosomes. We inferred this region to be the putative pseudoautosomal region (PAR) that is shared between the Z and W chromosome. The W scaffold measured 19.5 Mb in length and when the PAR is added to the length of the single W-assigned scaffold, the W is 78% of the predicted size as measured from karyotypes. The autosome-assigned scaffolds ranged in size from 195 to 65 Mb and differed from predicted lengths by a mean of 8.7% (range: 0.49-21.4%).

We validated the coverage-based assignments and the integrity of the fourteen large scaffolds by comparing heterozygosity and repetitive sequence content, performing PCR and qPCR analysis with markers designed from the genome assembly, isolating BAC clones, and using long-read sequencing to confirm scaffold continuity. Sex chromosomes should have lower levels of heterozygosity due to lower effective population sizes compared to autosomes, and single copy Z heterozygosity was significantly lower than the mean autosomal heterozygosity for both the female and male (*p* < 0.001 for both, Supplementary Figure S1). Another typical consequence of reduced sex chromosome effective population size is an accumulation of more repetitive sequences than present in autosomes. The density of repeat elements on the W and Z chromosomes was 41.5% and 20.3% higher compared to mean autosomal repeat density, and all categories showed repeat representation from Z/W chromosomes that were significantly different from null expectation (chi-square tests, all *p* < 0.001; Supplementary Figure S1). However, these patterns varied among transposable element categories which may reflect different histories of transposon activity between sex chromosomes. For example, W repeat density was less than that of autosomes for SINEs and less than Z chromosome repeat density for DNA/hAT elements (Supplementary Figure S1).

To validate the W chromosome scaffold specifically, we first identified eight multi-exonic genes that were located on both the Z and W scaffolds. We then designed primers targeting the W-linked versions of these genes. These markers amplified almost exclusively in females (Supplementary Results, Supplementary Table S3) and did not amplify in males. Sanger sequencing of PCR products confirmed that the correct target sequences were amplified. Additionally, we sequenced 11 bacterial artificial chromosome (BAC) clones that targeted four genes. The BAC clones containing a subset of the female-specific markers above (*OGT* and *RLIM*) mapped exclusively to the scaffold we identified as the W chromosome (Figure 2A). To validate the Z chromosome scaffold specifically, we conducted qPCR experiments using 22 loci (10 predicted on autosomes and 12 predicted on the Z). The average female/male ratios were 1.07 for autosomal and 0.61 for Z-linked loci, which closely matched the expected 1.0 ratio for autosomes and 0.5 ratio of the Z chromosome (Supplementary Results). The BAC clone containing *MED12* mapped back to the one-copy region of the Z-linked scaffold.

**Figure 2.**
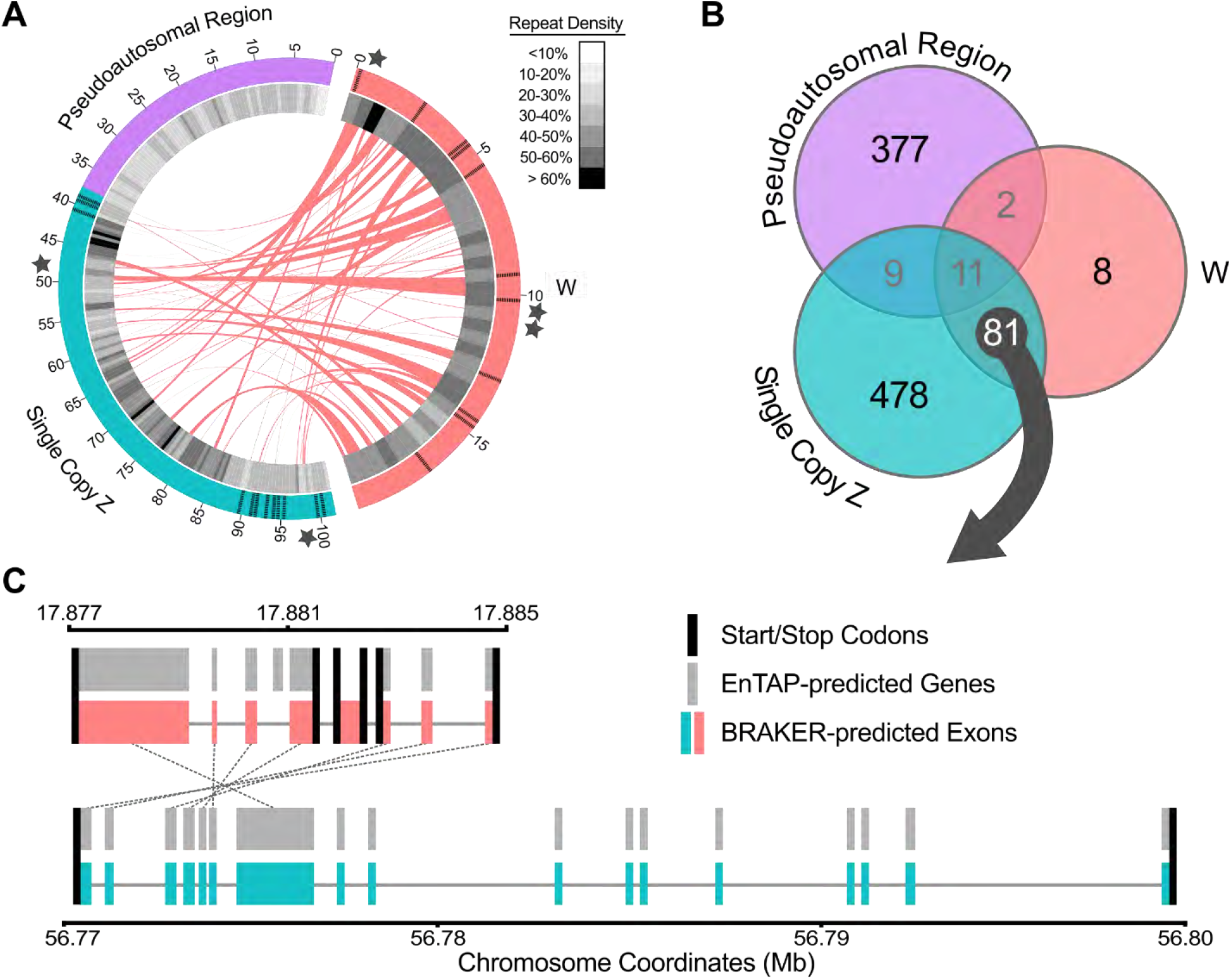
The W chromosome shares high sequence similarity with the Z chromosome, but is highly repetitive and functionally degraded. **(A)** The similarity between the W chromosome and the single copy region of the Z chromosome is visualized by a Circos^47^ plot for which links were established using one-to-one nucleotide alignments from NUCmer^48^. Each band represents at least one alignment greater than 500 bp in length, and multiple alignments are bundled together within 500 kb from each other. The inner ring displays the density of all repeat elements identified with RepeatModeler^49^ and RepeatMasker^50^ across 500 kb windows. The approximate location of validated W- and Z-linked PCR/qPCR (perpendicular hashed lines) and BAC clones (stars) are annotated on each scaffold. **(B)** A venn diagram shows the distribution of predicted gene models across the W, pseudoautosomal region, and the single copy Z. Listed of annotated protein-coding genes were extracted from EnTAP^46^ and cross referenced with coordinates from BRAKER^45^ annotation. When the 81 predicted genes that were identified in both W and single copy Z regions were compared, 38% display potential functional degradation in the copy located on the W. **(C)** An example of this pattern is the gene CAMSAP1. The copy located on the Z chromosome (bottom; teal boxes = exons from BRAKER, sum of gray boxes = predicted protein-coding exons from EnTAP, gray lines = introns) is bounded by a single start and stop codon (black bars) and each exon was identified by BRAKER and validated by EnTAP. In contrast, the copy located on the W is spread across three sets of start/stop codons, has shorter introns, shows disagreement between predicted exons and predicted proteins, and is missing several exons.

To validate continuity genome-wide, we collected long read sequencing for the same female used in the assembly. If the single copy nature of the Z/W chromosomes was responsible for undetected misjoins or assembly errors, these data would reflect a difficulty in aligning long reads to the Z/W compared to autosomes. We confirmed that the mapping to the Z/W chromosomes did not significantly differ in terms of coverage or the number of reads mapped when adjusted by scaffold size (Supplemental Table S4). These confirmatory PCR, qPCR, BAC sequencing, and long-read mapping experiments demonstrate that we have correctly assigned autosomal, Z-, and W-linked scaffolds. Collectively, these results illustrate that our genome assembly of the African Bullfrog provides chromosome-level resolution of all 12 autosomes, the Z/W sex chromosomes, and the subsequent ability to compare the evolution of sex-linked genes and sequences among vertebrates.

### Genetic divergence between the Z and W chromosome

Sex chromosomes typically share a common origin as evidenced by retained dosage sensitive genes and DNA sequence similarity, but their shared origin is also hidden by genomic divergence that accumulates during sex chromosome evolution^11,14,42–44^. To distinguish the genes that are shared or unique on the sex chromosomes of the African Bullfrog, we used RNA-Seq data from the same male and female individuals to predict gene models using BRAKER^45^ and identified 49,858 gene models. These gene models were filtered and annotated using EnTAP^46^ to provide a high confidence set of 14,716 protein-coding gene models: 13,656 for autosomes, 958 for the Z, and 102 for the W chromosome (Supplemental Table S5). The W chromosome had the lowest gene density and of the 102 genes only 8 were W specific. Instead, 79% (81/102) were also identified on the Z chromosome. To understand whether there was evidence of degradation of the gene copies on the W chromosome, we compared each of the 81 shared genes and found that 38% (31/81) had additional start/stop codons and changes to exon order and structure compared to the gene copy on the Z chromosome (Figure 2C). The relative lack of protein-coding genes, low gene density, asymmetry in gene structure, and changes to shared genes between the Z and W suggested a large degree of divergence between the Z and W chromosomes, which is consistent with typical models of sex chromosome evolution.

We then investigated if the differences in gene content between the Z, W, and autosomes reflected differences in gene function. We used Enrichr^51,52^ to generate and compare the gene ontology (GO^53^) terms of genes located on the Z (split into PAR and singly copy region), W, and autosomes. Genes annotated on the W were significantly enriched for Cellular Component GO terms associated with histone and heterochromatin function and Molecular Function GO terms associated with RNA binding (Hypergeometric tests with Benjamini-Hochberg *p* values < 0.5; Supplementary Table S6). Nine of the top ten enriched Biological Function GO terms for the W involved negative regulation of transcription (all *p* < 0.02). Significant enrichment of Cellular Component GO terms on the singly copy Z region was limited to microtubule organization (*p* = 0.04) and MHC class II protein synthesis (*p* = 0.002), while there was no significant enrichment among Biological Function GO terms. Both the W and single copy Z were enriched for multiple human phenotype ontologies^54^. For the W, these ontologies included X-linked recessive inheritance, abnormal salivation, and intellectual disability (all *p* < 0.002). For the Z, human phenotypes related to aggressive behavior, several anterior/cranial abnormalities (short nose, long face, short neck, mandibular prognathia), and X-linked dominant and recessive inheritance were enriched (all *p* < 0.001). Collectively, our analyses highlight that different gene functions are enriched on the Z compared to the W chromosome, and these differences make potential connections to the morphological and behavioral sexual dimorphism observed between African Bullfrog males and females^55^ while suggesting hypotheses for the homology of these sex-linked sequences.

### A Shared Origin of Genes Present On Vertebrate Heteromorphic Sex Chromosomes

To discover what sequences become heteromorphic on the sex chromosomes in the African bullfrog and to understand how these sequences vary among the evolutionary history of vertebrates, we aligned the African Bullfrog assembly to the genomes of the most well-assembled and annotated representative of five major vertebrates classes (human, chicken, green anole, African clawed frog, and half-smooth tongue sole fish) using PROmer^48^. The Z/W of African Bullfrogs shared significant similarity to large blocks of the ancestral human X chromosome and smaller alignments with the sole Z (Figure 3). Thirteen autosomal chromosomes had a greater number of alignments than expected by chance, including broad homology between the African Bullfrog Z and the African clawed frog 8, chicken 17 and 4, and the distal tip of human chromosome 9 (Figure 3). The extensive similarity of bullfrog sex chromosomes to the mammalian X and lesser similarity to the fish Z chromosome suggested either a common history of genomic sequences that become sex chromosomes or repeated use of the same sequences during sex chromosome evolution in vertebrates^56^.

**Figure 3.**
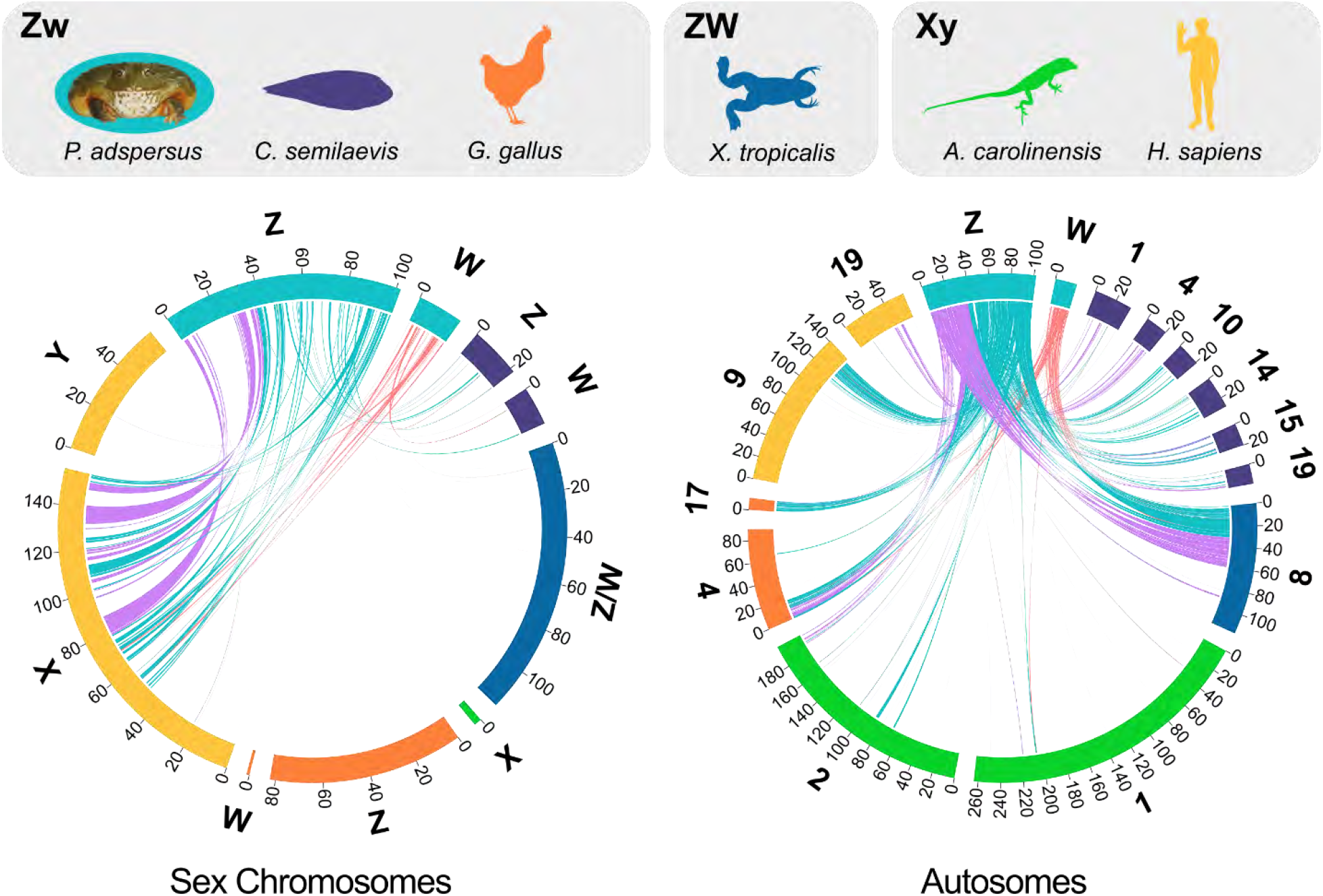
Comparison of vertebrate sex chromosomes to those of the African Bullfrog. The Z and W chromosome are homologous to large portions of the human X chromosome (left), but shows little similarity to other sex chromosomes. In contrast, the Bullfrog Z is homologous to the distal tip of human chromosome 9, portions of chromosome 8 of the African clawed frog, chicken 17 and 4, and multiple autosomes in the half-smooth tongue sole (right). Links between chromosomes were established using PROmer^48,57^ alignments that were filtered to only include unique query sequences, those greater than 100 bp, and at least 55% similarity. The lines originating from the Pyxicephalus Z/W represent areas where at least two sequences align within 1 kb from one another, and the lines are colored by assignment to the W (red), pseudoautosomal region (purple), or single copy Z region (blue). Only chromosomes with significant enrichment of putative orthologs are shown, where the total length of alignments was greater than what would be expected if the total number of alignments were distributed equally across all chromosomes.

According to the most well-supported model of sex chromosome evolution^8,42^, all vertebrate sex chromosomes—whether they have evolved convergently between divergent groups or share a common evolutionary history—should contain the genetic triggers for sex determination. More specifically, genes involved in the vertebrate sex determination network and their chromosome locations are the limited set of candidates that can canalize which genomic regions become heteromorphic sex chromosomes. To understand how homologous sequences between African Bullfrog sex chromosomes and other vertebrate chromosomes reflected the presence of shared genes related to sex determination, we curated a list of 67 genes involved in genetic sex determination^58^ (abbreviated as SDGs henceforth) or within the sex determination pathway^59^ of vertebrates and found the location of each gene in all six vertebrate genomes (Supplementary Table S7). In total, 21% (14/67) of these genes were located on one or more species’ sex chromosomes. The human X and African Bullfrog Z shared three of these genes: *SOX3* (the progenitor of the sex determination gene *SRY* in therian mammals), *ATRX* (found on the human X, the African Bullfrog Z, and in a degraded capacity on the African Bullfrog W), and *AR* (an androgen receptor gene; Supplemental Table S7). The Z chromosome of the smooth-tongue sole contained the greatest number of SDGs (7: *FST, DMRT1, DMRT3, Dapk1, Col27a1, Pcsk6*, and *Ereg*). The remaining SDGs that were physically located on heteromorphic sex chromosomes (N = 11) are limited to the sole, African clawed frog, African Bullfrog, and chicken. Surprisingly, no sex determination genes were found on the X chromosome of anole^22,60^. This observation may be due to lack of continuity in the anole assembly or, potentially, due to novel mechanisms of sex determination that have been recently discovered in other lizards^61^.

While the distribution of SDGs across heteromorphic sex chromosomes potentially reflects the diversity of landmarks initiating heteromorphic sex chromosome evolution across vertebrates^7,28,30^, these genes could share a common syntenic history that instead reflects functional characteristics associated with sex chromosome evolution. To address this possibility, we assigned each of the 67 SDGs to one of the ten reconstructed ancestral chromosomes of a putative vertebrate ancestor prior to the whole genome duplication event at the base of vertebrates^62^. While the SDGs were dispersed across eight of the ten ancestral chromosomes, the subset found on modern sex chromosomes were significantly more likely to be present on only two ancestral chromosomes (chi-square goodness of fit, *X^2^*_df=8_ = 20.67, *p* < 0.01) with 79% (11/14) of SDGs localized to ancestral chromosomes A and F (Figure 4). Ancestral chromosome A contained seven SDGs found on modern sex chromosomes that are split between the sole (N = 6), chicken (N = 4), African Bullfrog (N = 2), and African clawed frog (N = 1). The other ancestral chromosome (F) with more SDGs than expected contained four SDGs, three of which were shared between only humans and African Bullfrog. Furthermore, the distribution of these SDGs found on vertebrate sex chromosomes were not randomly distributed across the conserved vertebrate linkage (CVL) blocks that define the ancestral chromosomes and possess at least one SDG (8 of 41 CVLs, *X^2^*_df=40_ = 56.29, *p* = 0.02). Thus, while sex determining chromosomes and genes are labile across vertebrates, large chromosomal segments have been stable for more than 450 million years and only a restricted set of ancestral chromosomal blocks tend to become linked to heteromorphic sex chromosomes. In the unique case of humans and African Bullfrogs, both the human X and Bullfrog Z share an entire block of the ancestral vertebrate F chromosome after it underwent whole genome duplication prior to the evolution of jawed vertebrates (the Gnathostome ancestral chromosome F4^62^, Supplemental Table S7). These results suggest that not only are there repeatedly used sex determination trigger genes, but that two ancestral blocks of genomic sequences—and the genes they contain—are overwhelming represented among the dosage compromised sex chromosomes of vertebrates.

**Figure 4.**
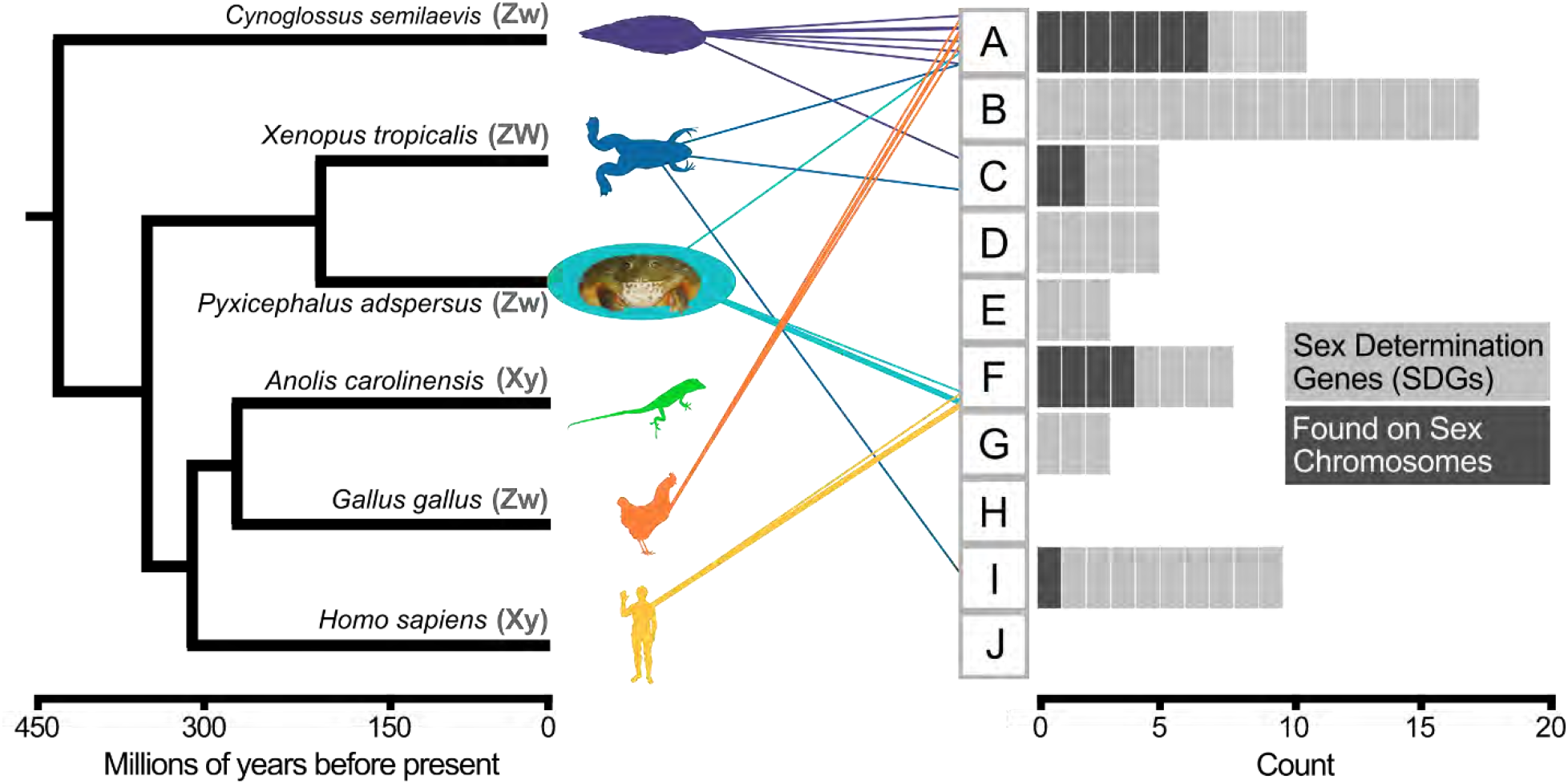
The sex chromosomes of Pyxicephalus adspersus fill a previous gap between the heteromorphic sex chromosomes identified in fish^63^ and those in reptiles^22,60^, birds^64,65^, and mammals^65^ ( dates from timetree.org^66,67^). The gray blocks (A-J) represent Nakatani et al.’s^62^ reconstruction of ten proto-vertebrate chromosome sequences. Sex determination genes (SDGs) that are located on the sex chromosomes of major vertebrate lineages can primarily be traced back to ancestral chromosome F for humans and A for all other lineages. However, African Bullfrogs show connections to both of these ancestral blocks.

### Shared ancestral blocks differ in haploinsufficiency tolerance

One of the most important problems associated with sex chromosome evolution is the reduction from two-copy to one-copy that occurs as sex chromosome pairs diverge in genetic content. Reductions in gene dosage are generally deleterious^16^ and the evolution of dosage compensation systems may help to rescue lineages from these effects^11,14,56^. That merely two ancestral genomic blocks make up the majority of modern vertebrate sex chromosomes suggests the possibility that there are variations in dosage sensitivity tied to these ancestral blocks that may constrain functional responses to dose during sex chromosome evolution.

Because the occurrence of A and F ancestral blocks on sex chromosomes corresponds to either lineages without clear signals of global dosage compensation (ancestral chromosome A: sole, clawed frog, chicken)^63,68^ and a lineage with global dosage compensation (ancestral chromosome F: human^69,70^), we asked whether genes found on these two ancestral chromosomes showed different dosage sensitivity characteristics. According to this hypothesis, genes on ancestral vertebrate chromosome F, which correspond to genes on the human X, should have increased sensitivity to changes in dosage compared to the genes found on ancestral chromosome A, which are found on heteromorphic sex chromosomes in species with no global dosage compensation^63,71^.

To test this hypothesis, we used an existing database of haploinsufficiency scores for the human genome and assigned scores to genes found on ancestral chromosome A and F in humans and the African Bullfrog. For both human and Bullfrog, genes on the sex chromosomes that map to ancestral chromosome F displayed increased haploinsufficiency on average compared to those that were linked with ancestral chromosome A (Figure 5). We similarly compared sensitivity to haploinsufficiency scores for genes that are retained as pairs on both heteromorphic sex chromosomes (X/Y pairs in humans and Z/W pairs in Bullfrogs). Increased sensitivity to haploinsufficiency across the few human genes that have X-Y pairs compared to those that remain as a single copy on the X is one of the fundamental lines of evidence for the necessary retention of certain genes on the Y^14^ and is also supported in ancestral Z-W paired genes in birds^11^. We replicated this result from humans and showed that the same pattern is mirrored in the 41 Bullfrog Z-W paired genes that have haploinsufficiency data available (Figure 5). Finally, to probe the hypothesis of dosage sensitivity further, we conducted functional enrichment analyses using Enrichr^52^ to examine human diseases associated with changes in dosage^72^. The set of genes shared on the sex chromosomes of humans and African Bullfrogs originating from the F block showed significant enrichment for 31 human diseases (Hypergeometric tests with Benjamini-Hochberg *p* values < 0.5). The majority of these diseases were related to both intellectual disability and X-linked recessive inheritance, such as Hunter, Partington, Allan-Herndon-Dudley, Opitz–Kaveggia, Fragile X, and Opitz-G/BBB syndromes. In contrast, the genes on the Bullfrog Z that originated from ancestral chromosome block A had no significant association with human diseases.

**Figure 5.**
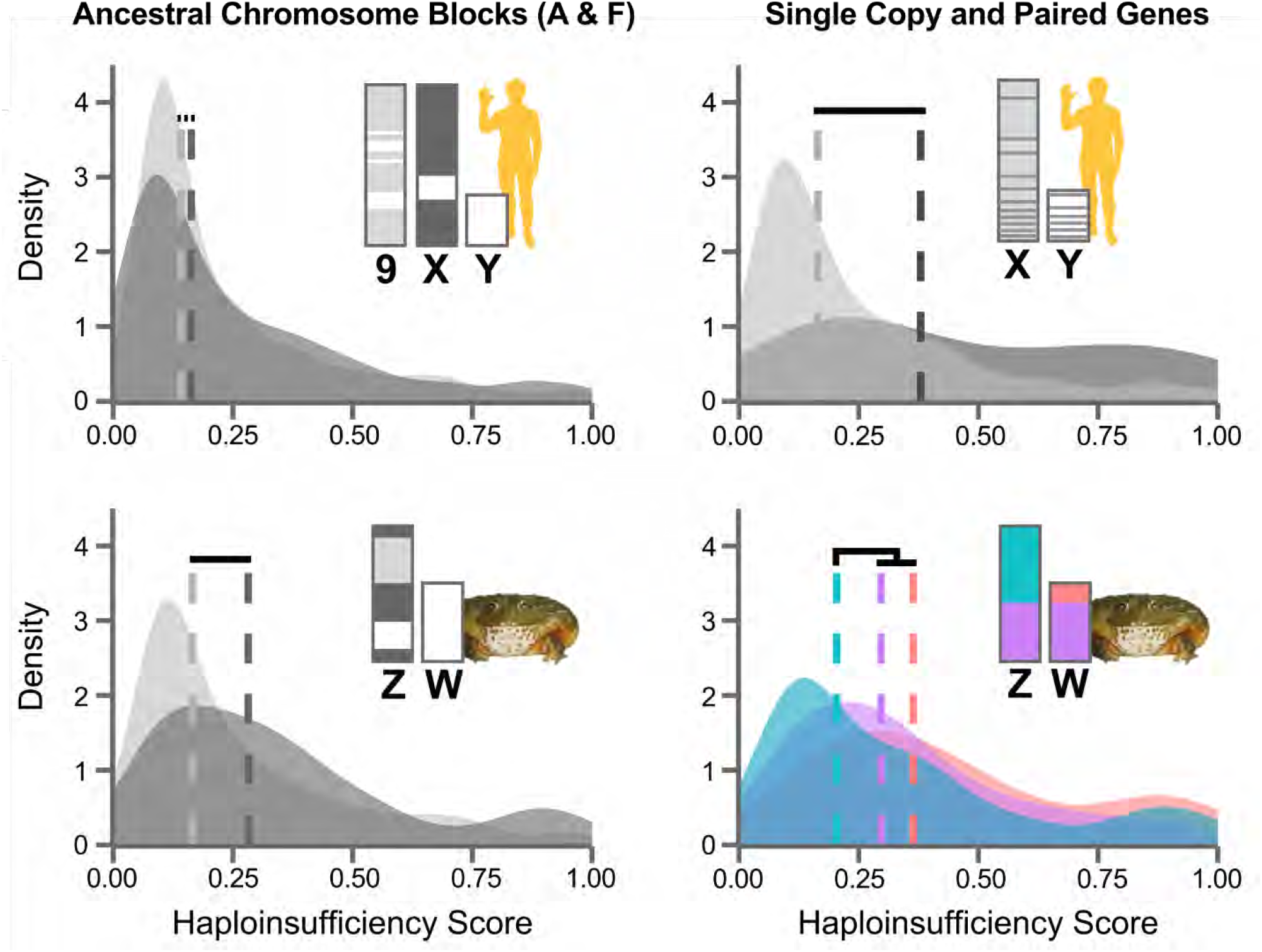
Haploinsufficiency analysis of human and frog suggests constraints on sequences that become sex chromosomes. In humans (top row) and African Bullfrogs (bottom row), the chromosome blocks that map to vertebrate ancestral chromosomes A and F differ in haploinsufficiency tolerance (left column) and genes that retain paired copies on both heteromorphic sex chromosomes have greater sensitivity to haploinsufficiency (right). The regions of human chromosomes that are associated with vertebrate ancestor chromosomes A (dark grey) and F (light grey) were delineated from Nakatani et al. (2007) and the genes present within these coordinates were assigned scores for haploinsufficiency tolerance^75^ (top left; N = 696 for A genes, N = 485 for F genes). We then identified those genes within the regions of the Bullfrog Z/W that aligned with human A/F, had the same gene name, and were also found in the human haploinsufficiency prediction database (bottom left; N = 218 for A genes, N = 225 for F genes). Dotted lines indicate the medians for distributions and the horizontal bars above lines are the results of two sample, one-directional Kolmogorov-Smirnov tests with p values < 0.05 (black lines) or < 0.10 (dotted line, top left). The right column subsets the genes on the human X/Y and the Bullfrog Z/W into those genes that share a copy between each sex chromosome (dark grey in humans and purple/red in Bullfrogs) and those that are at a single copy state on either the X (light grey) or Z (blue). Paired X-Y genes (N = 15) had significantly greater sensitivity to haploinsufficiency compared to unpaired genes (N = 699), matching previous analyses of the same data^14^. Likewise, genes located on the Bullfrog pseudoautosomal region (purple, N = 82) or with a copy on both the single copy Z and W chromosomes (N = 41) both had significantly greater sensitivity to haploinsufficiency compared to genes only present on the single copy region of the Z (N = 167, bottom right).

These results suggest that the two ancestral genomic blocks, which are repeatedly found on heteromorphic sex chromosomes in all vertebrates, have fundamentally different landscapes of dosage sensitivity that are consistent with the presence or absence of global dosage compensation systems. The sensitivity of genes associated with genes on ancestral chromosome F and now found on the mammalian X reinforces the hypothesis that dosage compensation through *XIST* as well as microRNA-mediated regulation may have been necessary to deal with the inherent dosage sensitivity of genes on the X as the Y chromosome diverged during sex chromosome evolution^73^. Conversely, genes from ancestral block A, which are found in sex chromosomes of fish, frogs, and birds, have less sensitivity to reductions in gene dosage and are consistent with the lack of global dosage compensation mechanisms in these organisms^63,68,74^. Taken together, the inherent difference in sensitivity of genes between the two ancestral genomic blocks that are found in nearly all vertebrate heteromorphic sex chromosomes may be a main driver of widespread functional responses to sex chromosome evolution in vertebrates.

## DISCUSSION

Homology across the sex chromosomes of major vertebrate groups was recognized more than five decades ago^27,76^, but this homology does not support a shared evolutionary history^56,65,77^. Our discovery of large scale similarity between the African Bullfrog Z chromosome and the human X echoes similarities between other vertebrates, such as the sole fish and chicken Z^63^ or the monotreme mammalian X and bird Z^65,78^. While each of these groups evolved sex chromosomes from different autosomes^77^, these observations combined suggest that repeated use of the same genomic sequences could be common during vertebrate sex chromosome evolution^28,29^. Using the newly assembled African Bullfrog genome and Nakatani and colleagues’ reconstruction of ancestral vertebrate chromosomes^62^, we showed that an overwhelming majority of genes that are associated with sex determination and located on vertebrate sex chromosomes are located on just two ancestral vertebrate chromosomes (A and F; Figure 4). These results suggest that certain ancestral genomic blocks are more likely to become associated with heteromorphic sex chromosomes, and this likelihood is not necessarily influenced by enrichment of sex pathway genes. Repeated sex-linkage of particular ancestral blocks may favor the evolution of heteromorphic sex chromosomes and create gene dosage imbalance that is molded for tolerance over evolutionary time.

Importantly, the inclusion of heteromorphic sex chromosomes from an amphibian prevents the human X/Y from being an outlier in the comparison of ancestral chromosome location, as the African Bullfrog is the only taxon that shares the same sex determination genes on their sex heteromorphic chromosomes. These genes are constrained to vertebrate ancestral chromosome F, but the Bullfrog Z also contains a sequence block derived from ancestral chromosome A, which is the primary contributor to the sex chromosomes of the sole and chicken. Because the corresponding A block found on the Bullfrog Z is located in autosome 9 of humans, this allowed us to to show that genes from the F block are more sensitive to haploinsufficiency compared to genes from the A block across autosomal (human), heterogametic male (human X), and heterogametic female (Bullfrog Z) states. Additionally, African Bullfrogs, like humans, have increased dosage sensitivity for those genes that remain in a paired state across both sex chromosomes. Given that sequences located on sex chromosomes are repeatedly reduced to a one copy state, these results support the hypothesis that genes originating from vertebrate ancestral chromosome F may require dosage compensation adjustment even though not all sex pathway genes are equally likely to become the sex determination trigger. In contrast, ancestral chromosome A possesses the greatest number of SDGs present on sex chromosomes—including highly influential sex determination genes such as *DMRT1*—but shows a more general resistance to haploinsufficiency. Because the African Bullfrog is now the only recognized species with both A and F blocks found in their heteromorphic sex chromosomes, future studies are needed to determine whether block F genes that became the mammal X are fundamentally sensitive to perturbation and required the evolution of dosage compensation or sex-biased selection for these genes on the X required dosage compensation secondarily.

Unlike mammals and birds, where sex chromosome states have remained relatively static for hundreds of millions of years, the presence of many shared genes between the African Bullfrog Z and W, the signals of functional degradation on the W, and the relatively large size of the pseudoautosomal region all point towards a relatively recent divergence between the Z and W. Additionally, because the sex-linked chromosome in the African clawed frog (chromosome 7) displays little similarity to the African Bullfrog Z/W, different autosomes have become sex chromosomes in these two frog lineages. These frog genomes provide a snapshot into the diversity of sex chromosomes in amphibians, which are thought to have evolved sex chromosomes in at least 20 independent instances^79^ and which have diverse combinations of sex determination mode (X/Y, Z/W) and relative size (heteromorphic/homomorphic)^3,79^. Furthermore, amphibians lack other confounding characteristics that affect sex chromosome evolution, such as the broad hermaphroditism present in fish or environmental sex determination in reptiles^3^. For these reasons, we anticipate that further work on the African Bullfrog and related amphibian taxa will gain increasing attention as model opportunities for the study of sex chromosome evolution and for understanding the links between gene dose, dosage compensation, and phenotype.

## METHODS

### Animals

This study complied with all ethical guidelines for working with amphibian animals and all procedures used in this paper were approved under University of Connecticut Institutional Animal Care Committee protocol number A13-030.

### Karyotype Analysis

We injected African Bullfrogs with 0.3% colchicine intraperitoneally. After 16 hours, frogs were anesthetized with MS-222, sexed, and the long bones were dissected and flushed twice with 0.075 M KCl. We incubated the cells at room temperature for 55 minutes. After incubation, the solution was centrifuged at 1400 rpm for 8 minutes. We removed the KCl solution and fixed the cell pellet in 6-10 mls of fresh 3:1 methanol:acetic acid. We cleaned microscope slides with EtOH and rinsed with water. We then added three to six drops of fixed cell suspension on clean microscope slides and allowed the sample to dry. Slides were added to Coplin jars containing Giemsa stain (1 Giemsa:18 Gurr buffer (ph 6.8)) with 50 mL dH_2_O for 2.5 minutes then rinsed in clean dH_2_O to remove excess stain. Slides were air-dried then sealed with mounting media and a coverslip. Finally, we examined chromosome spreads and photographed using an Olympus microscope.

### DNA Fragment Libraries and Short-Read Sequencing

We extracted high molecular weight DNA from blood from the same female and male examined for karyotypes using a QIAGEN DNAeasy blood and tissue kit (Hilden, Germany). We removed RNA by digestion with RNAase A. We evaluated DNA quality by gel electrophoresis and/or Bioanalyzer analysis. For short insert fragment libraries, We fragmented DNA to ~180 and 550 bp using Covaris sonication and prepared genomic libraries using an Illumina TruSeq PCR-free kit. For longer insert libraries (3, 6, and 12 kb), we ran DNA extractions on 1% TAE gels, evaluated size against NEB 2-log (Ipswich, MA) and Thermo Fisher HMW (Waltham, MA) DNA ladders, cut bands, and gel extracted DNA using a QIAGEN Gel Extraction kit at corresponding size ranges. We circularized, sheared, repaired, and amplified products to create mate-pair genomic libraries following protocols and reagents in the Illumina Nextera Mate Pair Library kit. We evaluated library complexity and concentration using a KAPPA Biosystems Library Quantification kit (Wilmington, Massachusetts, USA) and qPCR reactions. We prepared libraries for sequencing following standard protocols on the Illumina HiSeq 2000 to generate 2 x 100 bp reads. The 180 bp library was sequenced on three lanes to generate high coverage and all other libraries were sequenced on a single lane of Illumina flowcells.

For the female, larger 20-kb insert, Chicago, and HiC libraries were prepared. For the 20-kb library, DNA nuclei were extracted from oviduct tissue by homogenization with a glass dounce in 1X Dulbeccos Phosphate Buffered Saline. The lysate was filtered in miracloth and washed with buffer to release trapped nuclei. Nuclei were pelleted and placed into agarose using Bio-Rad (Herculus, CA) plug molds. Nuclei were extracted from gel plugs using GELase, and ~20 kb fragment lengths were cut from the gel and extracted using an EluTrap system. The 20-kb library was prepared by Lucigen (Middleton, WI) following protocols for the NxSeq Long Mate Pair Library Kit and was sequenced in two separate runs of a Illumina MiSeq with 2 x 250 bp read chemistry.

Three Chicago libraries were prepared as described previously^37^. Briefly, for each library, ~500 ng of HMW gDNA (mean fragment length = 100 kbp) was extracted from frozen heart tissue and the gDNA was reconstituted into chromatin *in vitro* and fixed with formaldehyde. The fixed chromatin was digested with DpnII, the 5’ overhangs were filled in with biotinylated nucleotides, and then free blunt ends were ligated. After ligation, crosslinks were reversed and the DNA was purified from protein. The purified DNA was treated to remove biotin that was not internal to ligated fragments. The DNA was then sheared to ~350 bp mean fragment size and sequencing libraries were generated using NEBNext Ultra enzymes and Illumina-compatible adapters. Finally, biotin-containing fragments were isolated using streptavidin beads before PCR enrichment of each library.

Two Dovetail Hi-C libraries were prepared from frozen liver tissue in a similar manner as described previously^81^. Briefly, for each library, chromatin in the nucleus was fixed with formaldehyde and then extracted. The fixed chromatin was digested with DpnII, the 5’ overhangs filled in with biotinylated nucleotides, and then free blunt ends were ligated. After ligation, formaldehyde cross links were reversed and the DNA was purified from protein. Purified DNA was treated to remove biotin that was not internal to ligated fragments. The DNA was sheared to ~350 bp mean fragment size and sequencing libraries were generated using NEBNext Ultra enzymes and Illumina-compatible adapters. Biotin-containing fragments were isolated using streptavidin beads before PCR enrichment of each library. Chicago and HiC libraries were sequenced on an Illumina HiSeq X to generate 2 x 150 bp reads.

### Read Processing

We assessed read qualities using the fastx-toolkit. We trimmed the first 8 bp of all 180 bp fragment library reads to remove nucleotide composition-biased bases. We removed putative contamination reads if they aligned to the NCBI environmental database by blastn (e-value < 1e-20) or if they aligned with MAPQ > 60 and > 80 bp alignment length to the human genome (Ensembl release 75) using BWA. We pre-processed the mate-pair data using NextClip v. 1.3^82^ to identify read pairs with junction adapter sequence in both mates (group A), first or second mate (group C and B, respectively), or mates without junction adapter sequence but sufficiently long reads after external adapter trimming (group D). We used Sickle (v. 1.33) to identify adaptors and trim reads from the 550 bp fragment library. For the 20 kb fragment library, we processed reads to identify true mates and to eliminate non-mate chimeras following custom scripts from Lucigen.

### Genome Assembly

We assembled the female sequencing data using ALLPATHS-LG, retaining contigs > 500 bp for the scaffolding steps. The ALLPATHS-LG assembly, shotgun reads, Chicago, and Dovetail Hi-C library reads were used as input data for HiRise, a software pipeline designed specifically for using proximity ligation data to scaffold genome assemblies^37^. An iterative analysis was conducted. First, shotgun and Chicago library sequences were aligned to the draft input assembly using a modified SNAP read mapper^83^. The separations of Chicago read pairs mapped within draft scaffolds were analyzed by HiRise to produce a likelihood model for genomic distance between read pairs, and the model was used to identify and break putative misjoins, to score prospective joins, and make joins above a threshold. After aligning and scaffolding Chicago data, Dovetail Hi-C library sequences were aligned and scaffolded following the same method. After scaffolding, shotgun sequences were used to close gaps between contigs to produce the final assembly. We identified one scaffold that matched to a bacteria sequence and six scaffolds that matched to mtDNA and removed these from the genome assembly. Assembly statistics of the ALLPATHS-LG input assembly and final HiRise assembly are provided in Supplementary Table S1. We identified repeat families using RepeatModeler^49^ with default parameters, and we produced a repeat-masked version of the African Bullfrog genome using RepeatMasker^50^.

### Sex-linked Scaffold Identification

To identify Z- and W-linked scaffolds, we separately mapped the female and male paired-end reads from all sequencing libraries to the repeat-masked^49,50^ female assembly—which contains both Z and W—using BWA-MEM^84^ 0.7.5. Next, we used the bedtools^85^ 2.27.1 *genomecov* function to calculate per-base sequencing depth of the mapped reads across all scaffolds (2.02 x 10^9^ and 2.07 x 10^9^ mapped reads for the female and male individual, respectively). We assigned each scaffold to either a Z chromosome, W chromosome, autosome (A), or unknown origin (U) based on the ratio between the male and female average depth. Given the number of A, Z, and W chromosomes in each sex, we expected autosomal scaffolds to have equal male and female coverage (1:1), Z scaffolds to have twice as much coverage from the male relative to the female (2:1), and W-linked scaffolds to have ~0 coverage from the male or ½-autosome coverage from the female. We repeated the calculation of depth using the samtools^86^ depth function to compare assignments from another method that also includes zero-coverage sites. All of the primary 14 scaffolds were clearly assigned to A, Z, or W (Figure 1), and 21 of the remaining 5,397 scaffolds with mismatched assignments between the two methods were identified by hand. In total, we were able to assign 93% of scaffolds, encompassing 97% of the total genome sequence to autosomes, Z, or W, with the remainder unassigned because of ambiguity in the male-female comparisons. We inferred synteny between the putative W and Z chromosomes using a NUCmer^87^ (v3.23) alignment that was filtered using both the -q and -r flag (one-to-one alignments).

### Repeat Content and Heterozygosity

The output of RepeatModeler/RepeatMasker^49,50^ 4.0.6 was parsed using the R^41^ package *dplyr* and the number of repeat elements per scaffold size was calculated for the most-represented repeat families. For visualizing repeat content of the Z and W chromosomes, we divided the Bullfrog scaffolds into 500 kb windows using bedtools^85^ *makewindows* and then used bedtools *intersect* with the windows and the coordinates produced by RepeatMasker. We calculated heterozygosity using the GATK^88^ *HaplotypeCaller*, converted the resultant VCF to a table, and calculated the number of INDEL and SNP heterozygous sites divided by the size of each scaffold.

### W- and Z-linked locus development

To validate W chromosome assignments, we designed primers to match sequence locations in our assembly. For W-linked loci, we searched gene models for coding regions, extracted sequences, then used Primer3 v 2.3.7^89^ to select primer locations that would generate amplicons 350-700 bases in length. Amplicons were blasted against the genome assembly and the results were filtered to generate the best primer pair. We designed primers for eight alleged W-linked loci (*OGT, THOC2, NLGN3, FOXO4, RLIM, DLG3, ZMYM3*, and *PRPS1*), a locus that matched both an autosomal and W-linked scaffold (*AMMECR1*), and a locus that matched W- and Z-linked scaffolds (*ZC3H12B*). We extracted DNA from blood tissue from 8 additional female and 12 additional male *Pyxicephalus adspersus* using a QIAGEN DNeasy blood and tissue kit (Hilden, Germany) and conducted PCR reactions followed by gel visualization to confirm amplification for the W-specific markers (full reaction conditions in supplemental methods).

To validate Z and autosome assignments, we designed primers to match coding sequences that were computationally identified in scaffolds and to conduct qPCR experiments to check copy number. Primers were identified to target the Z chromosome using similar methods as the W-loci, and amplicons were blasted against the genome assembly and filtered for the best primer pair. We designed qPCR primers for 10 autosomal loci (*BMP5, DMRT1, DMRT2, DNAJC11, EEF1A1, FOXL2, NGLY1, SOX3, SOX9*, and *WNT4*) and 12 alleged Z-linked loci (*ALG13, APEX2, AR, ATG4A, CAPN6, FAM120A, GPR21, MED12, PAPPA, PCDH11X, STARD8*, and *STRBP*). A pair of primers for *ABL1* matched both an autosomal and Z-linked location. Quantitative PCR reactions were measured on a CFX96TM Optical Reaction Module thermocycler (Bio-Rad, California, USA; full reaction conditions in supplementary methods). To evaluate copy number differences between females and males, we performed qPCR with 8 eight female and eight male DNA samples and compared the normalized female/male Ct ratios.

### Bacterial Artificial Chromosome libraries

A 10X coverage BAC library was prepared by the Clemson University Genomics Institute using megabase size DNA extracted from oviduct tissue from the same female used for the genome assembly. Nuclei was isolated from oviduct tissue by gently macerating with a razor blade and breaking apart the tissue using a dounce loose fitting pestle (Chemglass Life Sciences, Vineland, New Jersey, USA) in 1% phosphate buffered saline (PBS) with 0.5 M EDTA, ph 8.0. The homogenzied tissue was filtered through one layer of miracloth (Millipore, Billerica, MA) into a 50-ml conical tube, and the total volume was adjusted to 45 ml with PBS and nuclei-pelleted by centrifugation (1800 g, 4°C) using a swinging-bucket rotor. The nuclei were washed two additional times with PBS and gently re-suspended in 1 ml of PBS before being warmed to 50°C. Nuclei plugs were prepared by adding 1 ml of 1.5% low-melt agarose and aliquoted into plug molds (Bio-Rad, Hercules, California).

Two independent restriction-derived BAC libraries were prepared by digesting the megabase-size DNA with HindIII and BstYI restriction enzymes to maximize the coverage of the genome with long inserts. Size selected DNA (>150 kb) was ligated with cut/linearized and dephosphorylated BAC vector and then electroporated into *E. coli* DH10B electrocompetent cells. BAC transformants were selected on LB plates containing chloramphenicol, X-Gal and IPTG. White recombinant colonies were picked robotically using the Genetix Q-bot and individual clones were stored in Genetix 384-well microtiter plates as glycerol stocks at −80°C. The library was evaluated for insert size by sampling 384 clones at random, miniprepping, releasing the inserts with NotI, and running the DNA on CHEF gels. The library had inserts on average of at least 120 kb and <5% had empty vectors or non-recombinants. The BAC library was plated on high-density colony filters using a Genetix Q-bot in 4×4 double-spotted arrays (18,432 clones represented per filter). Filters were arrayed on GE HealthCare Hybond N+. This library macroarray was used to query homoeologous BACs for sequencing and the library is available upon request.

Overgo probes were designed based on PCR amplicons of four gene models that we hypothesized to be autosomal (*DMRT1*) or sex-linked (W: *OGT* and *RLIM*; Z: *MED12*). BACs identified by overgo hybridization were picked, grown, and plated in LB media^90^. BAC clones were cut and sequenced these using Pacific Biosciences RS II and P5/C3 chemistry for long read sequencing. Raw reads were corrected and assembled using CANU^91^. The resulting contigs were screened for vector bases with CrossMatch and Sanger BAC-end sequences were added and final manual assemblies were performed using Consed^92^. BAC annotations were performed with MAKER-P^93,94^ (v. 2.31.8) to identify gene models and homologous proteins were identified with tbalstx. We mapped sequences to the genome assembly using BLAT^95^ v36.2.

### Additional long read sequencing

We gathered long read sequencing data from the same female used for the genome assembly at the New York Genome Center and the SMRT Sequencing Lab at Weill Cornell Medical College. High molecular weight DNA was extracted from blood tissue and the 20 kb Blue Pippin version of the PacBio library preparation protocol and used with the P5 Enzyme in conjunction with C3 chemistry. We sequenced five SMRT cells on a PacBio RS II for 180 minutes per SMRT cell and data were filtered using SMRT Analysis v 2.2. Reads were corrected using LSC^96^ and NUCmer^48^ alignments to a preliminary ALLPATHS-LG assembly of the African Bullfrog genome. After correction, 206,654 sequences (mean length = 5.48 kb) were mapped to the current African Bullfrog assembly using blasr^97^.

### Genome Alignments

We aligned the repeat-masked Bullfrog genome to the genomes of human (v38, Ensembl), chicken (v5, Ensembl), *Anolis carolinensis* (v2.0, Ensembl), *Xenopus tropicalis* (assembly version 9.1, www.xenbase.org), and half-smooth tongue sole (version 1.0, NCBI, GenBank assembly ID GCA_000523025.1) using PROmer v3.23^87^. The resultant delta files were filtered with the -q flag to ensure only unique query alignments remained, and any alignments < 100 bp long or < 55% similar were filtered out.

### Transcriptome Libraries

We extracted total RNA from male testis and female brain tissue using Trizol and a BeadBug homogenizer (Benchmark Scientific, Sayreville, New Jersey, USA). We evaluated RNA quality on a Bioanalyer. We then built stranded mRNA transcriptome libraries for each tissue following standard procedures of an Illumina TruSeq Stranded mRNA Library kit. Finally, we sequenced libraries on an Illumina MiSeq instrument using 2 x 75 bp read chemistry.

### Annotation and Ortholog Identification

We predicted gene models for the African Bullfrog assembly using BRAKER^45^ v1.9. First, we removed ambiguities from the assembly by resolving each ambiguity code as one of the potential bases at random. We then used TopHat2^98^ to map a modest RNA-seq data set (14 million reads) collected from the same male and female individual as above to the genome assembly. The results from BRAKER were parsed using a custom script to compile basic statistics. The protein models produced by BRAKER were then annotated using the Eukaryotic Non-Model Transcriptome Annotation Pipeline (EnTAP^46^ v0.8), which conducts similarity searches, filters for contamination, and assigns proteins to families and ontology groups. We included UniProt^99^, RefSeq^100^, and NCBI non-redundant protein databases in our similarity search, filtered out matches to fungal and bacterial taxa, and used taxonomic ‘favoring’ of hits. EnTAP output was filtered for only those sequences that had predicted proteins from EggNOG-Emapper^101,102^ v0.7.4.1 and then combined with genome coordinates from BRAKER to provide the scaffold locations for the predicted protein-coding genes. Finally, we assembled four lists of predicted protein-coding genes according to their genomic location (autosomal, W, pseudoautosomal region, and singly copy Z region) and measured functional enrichment using hypergeometric test with Benjamini-Hochberg corrected *p* values as implemented by Enrichr^51,52^.

### Synteny of sex-linked sequences in vertebrates

We curated a list of genes in vertebrates that are directly linked to sex determination or within sex determination pathways (Supplementary Table S7; 32 from Capel (2017)^59^, 29 from Forconi *et al*. (2013)^58^, and 6 additional). This list was used to located each gene in the genome of all six taxa using a combination of direct searches via NCBI and cross referencing with the previous PROmer alignments. To ask if genes on this list that are located on heteromorphic sex chromosomes are overrepresented on ancestral chromosomes or blocks, we used the locations on the human genome to assign each gene an ancestral location based on Nakatani *et al*’s (2007)^62^ reconstruction of the pre-duplicated chromosome set of a putative vertebrate ancestor. We tested if the distribution of ancestral chromosome locations was statistically different between those sex-determination genes located on sex chromosomes and those that were located in autosomes using a Chi-square goodness of fit test.

### Quantification of Haploinsufficiency

First, we downloaded the database of human gene-by-gene haploinsufficiency scores from Huang *et al*. (2010) and extracted from this list those genes that fell within areas delineated by Nakatani *et al* (2007) as part of ancestral vertebrate chromosome F (majority of human X, N = 484) and ancestral vertebrate chromosome A (human 9, N = 696). While portions of other human chromosomes also have blocks that map to A, we chose to only use those on chromosome 9 because 1) this chromosome allows for less ambiguity in ancestral assignment due to A being the only representative ancestral block present and 2) chromosome 9 is the autosomal location of the majority of A block genes associated with the sex chromosomes of the sole and chicken. These lists of genes and corresponding haploinsufficiency scores were then cross referenced with the lists of genes on the Bullfrog Z/W and their coordinates (pseudoautosomal, singly copy Z, W). Finally, we extracted the scores for the 15 ancestrally paired X-Y genes curated by Bellott *et al*. (2014) and compared their haploinsufficiency scores to the scores for the remaining single copy genes on the X (N = 699). We quantified differences in haploinsufficiency score between A and F blocks or paired/non-paired genes using two-sample Kolmogorov-Smirnov tests in R^41^. Because we predicted that F block and paired genes would have greater haploinsufficiency sensitivity when compared to A block and unpaired genes, respectively, we used an alternative hypothesis of “greater” instead of a two-sided alternative.

## ACKNOWLEDGEMENTS

We thank Rachel O’Neill and Judy Brown for providing use of microscopes and karyotype advice, Bo Reece and the UConn Center for Genome Innovation for advice and support, Dayna Oschwald and Noah Alexander from the New York Genome Center and Weill Cornell Medical College SMRT Sequencing Lab for facilitating and performing PacBio sequencing, Sierra Wilson from Dovetail Genomics for leading Chicago and HiC library sequencing, Brenton Graveley for access to the Illumina MiSeq instrument, Christopher Saski from the Clemson University Genomics Institute for BAC library construction, Jill Wegrzyn, Sumaira Zaman, Neranjan V. Perera, and the UConn Bioinformatics Core for advice and help with gene annotation, Andrew Galinski, Pedro Cuc-Saenz, and Andrew Villion for help with PCR and qPCR experiments, and Hannah Ralicki for help with karyotype and computational analysis. Funding for this project was provided by UConn startup funds and NSF XSEDE research allocation awards TG-DMS140018 and TG-MCB141026 to John Malone.

## AUTHOR CONTRIBUTIONS

RDD analyzed data, designed figures, and wrote the manuscript. RSK constructed genomic libraries and helped perform karyotype experiments. JWM developed code and analyzed preliminary datasets. LDP provided logistical and intellectual support. JHM performed computational analyses, wrote the manuscript, and directed the project. All authors have reviewed and approved the manuscript.

## SUPPLEMENTARY INFORMATION

Supplementary Data contains the following individual files:

*Data_Padspersus_Annotated_Genes*
*Supplementary Material File*
*Supplementary Methods and Results*
*Figure S1_Repeat Element and Heterozygosity Comparisons*
*Table S1_Assembly Statistics*
*Table S2_Library Stats and Mapping Table*
*S3_PCR-qPCR-BAC*
*Table S4_PacBio Mapping*
*Table S5_Scaffold Assignments and Annotation Statistics*
*Table S6_Functional Enrichment Table*
*S7_Conserved Vertebrate Linkage*

## REFERENCES

1. Neiman, M., Lively, C. M. & Meirmans, S. Why sex? A pluralist approach revisited. Trends Ecol. Evol. 32, 589–600 (2017).

2. Bachtrog, D. et al. Sex determination: why so many ways of doing it? PLoS Biol. 12, e1001899 (2014).

3. Tree of Sex Consortium. Tree of Sex: a database of sexual systems. Sci Data 1, 140015 (2014).

4. Ellegren, H. Sex-chromosome evolution: recent progress and the influence of male and female heterogamety. Nat. Rev. Genet. 12, 157–166 (2011).

5. Pan, Q. et al. Vertebrate sex-determining genes play musical chairs. C. R. Biol. 339, 258–262 (2016).

6. Mank, J. E. Sex chromosome dosage compensation: definitely not for everyone. Trends Genet. 29, 677–683 (2013).

7. Bachtrog, D. et al. Are all sex chromosomes created equal? Trends Genet. 27, 350–357 (2011).

8. Bachtrog, D. Y-chromosome evolution: emerging insights into processes of Y-chromosome degeneration. Nat. Rev. Genet. 14, 113–124 (2013).

9. Bull, J. J. Evolution of Sex: Determining Mechanisms. (The Benjamin/Cummings Publishing Company Inc., 1983).

10. Hughes, J. F. & Page, D. C. The biology and evolution of mammalian Y chromosomes. Annu. Rev. Genet. 49, 507–527 (2015).

11. Bellott, D. W. et al. Avian W and mammalian Y chromosomes convergently retained dosage-sensitive regulators. Nat. Genet. 49, 387–394 (2017).

12. Schartl, M., Schmid, M. & Nanda, I. Dynamics of vertebrate sex chromosome evolution: from equal size to giants and dwarfs. Chromosoma (2015).

13. Cody, J. D. & Hale, D. E. Linking chromosome abnormality and copy number variation. Am. J. Med. Genet. A 155A, 469–475 (2011).

14. Bellott, D. W. et al. Mammalian Y chromosomes retain widely expressed dosage-sensitive regulators. Nature 508, 494–499 (2014).

15. Cortez, D. et al. Origins and functional evolution of Y chromosomes across mammals. Nature 508, 488–493 (2014).

16. Chen, Z.-X., Golovnina, K., Sultana, H., Kumar, S. & Oliver, B. Transcriptional effects of gene dose reduction. Biol. Sex Differ. 5, 5 (2014).

17. Birchler, J. A. & Veitia, R. A. Gene balance hypothesis: connecting issues of dosage sensitivity across biological disciplines. Proc. Natl. Acad. Sci. U. S. A. 109, 14746–14753 (2012).

18. Plath, K., Mlynarczyk-Evans, S., Nusinow, D. A. & Panning, B. Xist RNA and the mechanism of X 18 chromosome inactivation. Annu. Rev. Genet. 36, 233–278 (2002).

19. Kharchenko, P. V., Xi, R. & Park, P. J. Evidence for dosage compensation between the X chromosome and autosomes in mammals. Nat. Genet. 43, 1167–1169 (2011).

20. Lin, H. et al. Dosage compensation in the mouse balances up-regulation and silencing of X-linked genes. PLoS Biol. 5, e326 (2007).

21. Marin, R. et al. Convergent origination of a Drosophila-like dosage compensation mechanism in a reptile lineage. Genome Res. 27, 1974–1987 (2017).

22. Rupp, S. M. et al. Evolution of Dosage Compensation in *Anolis carolinensis*, a Reptile with XX/XY Chromosomal Sex Determination. Genome Biol. Evol. 9, 231–240 (2017).

23. Smith, G., Chen, Y.-R., Blissard, G. W. & Briscoe, A. D. Complete dosage compensation and sex-biased gene expression in the moth *Manduca sexta*. Genome Biol. Evol. 6, 526–537 (2014).

24. Gu, L. & Walters, J. R. Evolution of Sex Chromosome Dosage Compensation in Animals: A Beautiful Theory, Undermined by Facts and Bedeviled by Details. Genome Biol. Evol. 9, 2461–2476 (2017).

25. Vicoso, B., Emerson, J. J., Zektser, Y., Mahajan, S. & Bachtrog, D. Comparative sex chromosome genomics in snakes: differentiation, evolutionary strata, and lack of global dosage compensation. PLoS Biol. 11, e1001643 (2013).

26. Vicoso, B. & Bachtrog, D. Lack of global dosage compensation in Schistosoma mansoni, a female-heterogametic parasite. Genome Biol. Evol. 3, 230–235 (2011).

27. Ezaz, T., Stiglec, R., Veyrunes, F. & Marshall Graves, J. A. Relationships between vertebrate ZW and XY sex chromosome systems. Curr. Biol. 16, R736–43 (2006).

28. O’Meally, D., Ezaz, T., Georges, A., Sarre, S. D. & Graves, J. A. M. Are some chromosomes particularly good at sex? Insights from amniotes. Chromosome Res. 20, 7–19 (2012).

29. Marshall Graves, J. A. & Peichel, C. L. Are homologies in vertebrate sex determination due to shared ancestry or to limited options? Genome Biol. 11, 205 (2010).

30. Graves, J. A. M. Evolution of vertebrate sex chromosomes and dosage compensation. Nat. Rev. Genet. 17, 33–46 (2016).

31. Blaser, O., Grossen, C., Neuenschwander, S. & Perrin, N. Sex-chromosome turnovers induced by deleterious mutation load. Evolution 67, 635–645 (2013).

32. Gnerre, S. et al. High-quality draft assemblies of mammalian genomes from massively parallel sequence data. Proc. Natl. Acad. Sci. U. S. A. 108, 1513–1518 (2011).

33. Butler, J. et al. ALLPATHS: de novo assembly of whole-genome shotgun microreads. Genome Res. 18, 810–820 (2008).

34. Maccallum, I. et al. ALLPATHS 2: small genomes assembled accurately and with high continuity from short paired reads. Genome Biol. 10, R103 (2009).

35. Schmid, M. Chromosome Banding in Amphibia V. Highly Differentiated ZW/ZZ Sex Chromosomes and Exceptional Genome Size in *Pyxicephalus adspersus* (Anura, Ranidae). Chromosoma 80, 69–96 (1980).

36. Schmid, M. & Bachmann, K. A frog with highly evolved sex chromosomes. Experientia 37, 243–245 (1981).

37. Putnam, N. H. et al. Chromosome-scale shotgun assembly using an in vitro method for long-range linkage. Genome Res. 26, 342–350 (2016).

38. Simão, F. A., Waterhouse, R. M., Ioannidis, P., Kriventseva, E. V. & Zdobnov, E. M. BUSCO: assessing genome assembly and annotation completeness with single-copy orthologs. Bioinformatics 31, 3210–3212 (2015).

39. Hellsten, U. et al. The genome of the Western clawed frog *Xenopus tropicalis*. Science 328, 633–636 (2010).

40. Session, A. M. et al. Genome evolution in the allotetraploid frog *Xenopus laevis*. Nature 538, 336–343 (2016).

41. R Core Team. R: A language and environment for statistical computing. (R Foundation for Statistical Computing, 2017).

42. Charlesworth, D., Charlesworth, B. & Marais, G. Steps in the evolution of heteromorphic sex chromosomes. Heredity 95, 118–128 (2005).

43. Vicoso, B. & Charlesworth, B. Evolution on the X chromosome: unusual patterns and processes. Nat. Rev. Genet. 7, 645–653 (2006).

44. Zhou, Q. et al. Complex evolutionary trajectories of sex chromosomes across bird taxa. Science 346, 1246338 (2014).

45. Hoff, K. J., Lange, S., Lomsadze, A., Borodovsky, M. & Stanke, M. BRAKER1: Unsupervised RNA-Seq-based genome annotation with GeneMark-ET and AUGUSTUS. Bioinformatics 32, 767–769 (2016).

46. Hart, A. J. et al. EnTAP: Bringing faster and smarter functional annotation to non-model Eukaryotic transcriptomes. bioRxiv 307868 (2018). doi:10.1101/307868

47. Krzywinski, M. et al. Circos: an information aesthetic for comparative genomics. Genome Res. 19, 1639–1645 (2009).

48. Marçais, G. et al. MUMmer4: A fast and versatile genome alignment system. PLoS Comput. Biol. 14, e1005944 (2018).

49. Smit, A. & Hubley, R. RepeatModeler Open-1.0. (2008).

50. Smit, A., Hubley, R. & Green, P. RepeatMasker Open-4.0. (2013).

51. Chen, E. Y. et al. Enrichr: interactive and collaborative HTML5 gene list enrichment analysis tool. BMC Bioinformatics 14, 128 (2013).

52. Kuleshov, M. V. et al. Enrichr: a comprehensive gene set enrichment analysis web server 2016 update. Nucleic Acids Res. 44, W90–7 (2016).

53. Ashburner, M. et al. Gene ontology: tool for the unification of biology. The Gene Ontology Consortium. Nat. Genet. 25, 25–29 (2000).

54. Köhler, S. et al. The Human Phenotype Ontology in 2017. Nucleic Acids Res. 45, D865–D876 (2017).

55. Hayes, T. & Licht, P. Gonadal involvement in sexual size dimorphism in the African bullfrog (*Pyxicephalus adspersus*). J. Exp. Zool. 264, 130–135 (1992).

56. Ezaz, T., Srikulnath, K. & Graves, J. A. M. Origin of amniote sex chromosomes: an ancestral super-sex chromosome, or common requirements? J. Hered. 108, 94–105 (2017).

57. Delcher, A. L., Salzberg, S. L. & Phillippy, A. M. Using MUMmer to identify similar regions in large sequence sets. Curr. Protoc. Bioinformatics Chapter 10, Unit 10.3 (2003).

58. Forconi, M. et al. Characterization of sex determination and sex differentiation genes in *Latimeria*. PLoS One 8, e56006 (2013).

59. Capel, B. Vertebrate sex determination: evolutionary plasticity of a fundamental switch. Nat. Rev. Genet. 18, 675–689 (2017).

60. Alföldi, J. et al. The genome of the green anole lizard and a comparative analysis with birds and mammals. Nature 477, 587–591 (2011).

61. Holleley, C. E. et al. Sex reversal triggers the rapid transition from genetic to temperature-dependent sex. Nature 523, 79–82 (2015).

62. Nakatani, Y., Takeda, H., Kohara, Y. & Morishita, S. Reconstruction of the vertebrate ancestral genome reveals dynamic genome reorganization in early vertebrates. Genome Res. 17, 1254–1265 (2007).

63. Chen, S. et al. Whole-genome sequence of a flatfish provides insights into ZW sex chromosome evolution and adaptation to a benthic lifestyle. Nat. Genet. 46, 253–260 (2014).

64. International Chicken Genome Sequencing Consortium. Sequence and comparative analysis of the chicken genome provide unique perspectives on vertebrate evolution. Nature 432, 695–716 (2004).

65. Bellott, D. W. et al. Convergent evolution of chicken Z and human X chromosomes by expansion and gene acquisition. Nature 466, 612–616 (2010).

66. Hedges, S. B., Marin, J., Suleski, M., Paymer, M. & Kumar, S. Tree of life reveals clock-like speciation and diversification. Mol. Biol. Evol. 32, 835–845 (2015).

67. Kumar, S., Stecher, G., Suleski, M. & Hedges, S. B. TimeTree: a resource for timelines, timetrees, and divergence times. Mol. Biol. Evol. 34, 1812–1819 (2017).

68. Itoh, Y. et al. Dosage compensation is less effective in birds than in mammals. J. Biol. 6, 2 (2007).

69. Nguyen, D. K. & Disteche, C. M. Dosage compensation of the active X chromosome in mammals. Nat. Genet. 38, 47–53 (2006).

70. Disteche, C. M. Dosage compensation of the sex chromosomes. Annu. Rev. Genet. 46, 537–560 (2012).

71. Wang, Q., Mank, J. E., Li, J., Yang, N. & Qu, L. Allele-specific expression analysis does not support sex chromosome inactivation on the chicken Z chromosome. Genome Biol. Evol. 9, 619–626 (2017).

72. Pletscher-Frankild, S., Pallejà, A., Tsafou, K., Binder, J. X. & Jensen, L. J. DISEASES: text mining and data integration of disease-gene associations. Methods 74, 83–89 (2015).

73. Naqvi, S., Bellott, D. W., Lin, K. S. & Page, D. C. Conserved microRNA targeting reveals preexisting gene dosage sensitivities that shaped amniote sex chromosome evolution. Genome Res. (2018).

74. Malone, J. H. Dosage compensation in frogs and toads. in Epigenetics: Current Research and Emerging Trends (ed. Chadwick, B. P.) 167–183 (Caister Academic Press, 2015).

75. Huang, N., Lee, I., Marcotte, E. M. & Hurles, M. E. Characterising and predicting haploinsufficiency in the human genome. PLoS Genet. 6, e1001154 (2010).

76. Ohno, S. Sex Chromosomes and Sex-Linked Genes. (Springer-Verlag Berlin Heidelberg, 1967).

77. Matsubara, K. et al. Evidence for different origin of sex chromosomes in snakes, birds, and mammals and step-wise differentiation of snake sex chromosomes. Proc. Natl. Acad. Sci. U. S. A. 103, 18190–18195 (2006).

78. Veyrunes, F. et al. Bird-like sex chromosomes of platypus imply recent origin of mammal sex chromosomes. Genome Res. 18, 965–973 (2008).

79. Evans, B. J., Alexander Pyron, R. & Wiens, J. J. Polyploidization and sex chromosome evolution in amphibians. in Polyploidy and Genome Evolution 385–410 (Springer, Berlin, Heidelberg, 2012).

80. Malcom, J. W., Kudra, R. S. & Malone, J. H. The sex chromosomes of frogs: variability and tolerance offer clues to genome evolution and function. J Genomics 2, 68–76 (2014).

81. Lieberman-Aiden, E. et al. Comprehensive mapping of long-range interactions reveals folding principles of the human genome. Science 326, 289–293 (2009).

82. Leggett, R. M., Clavijo, B. J., Clissold, L., Clark, M. D. & Caccamo, M. NextClip: an analysis and read preparation tool for Nextera Long Mate Pair libraries. Bioinformatics 30, 566–568 (2014).

83. Zaharia, M. et al. Faster and More Accurate Sequence Alignment with SNAP. arXiv [cs.DS] (2011).

84. Li, H. Aligning sequence reads, clone sequences and assembly contigs with BWA-MEM. arXiv [q-bio.GN] (2013).

85. Quinlan, A. R. & Hall, I. M. BEDTools: a flexible suite of utilities for comparing genomic features. Bioinformatics 26, 841–842 (2010).

86. Li, H. A statistical framework for SNP calling, mutation discovery, association mapping and population genetical parameter estimation from sequencing data. Bioinformatics 27, 2987–2993 (2011).

87. Kurtz, S. et al. Versatile and open software for comparing large genomes. Genome Biol. 5, R12 (2004).

88. McKenna, A. et al. The Genome Analysis Toolkit: a MapReduce framework for analyzing next-generation DNA sequencing data. Genome Res. 20, 1297–1303 (2010).

89. Untergasser, A. et al. Primer3--new capabilities and interfaces. Nucleic Acids Res. 40, e115 (2012).

90. Saski, C. A., Feltus, F. A., Parida, L. & Haiminen, N. BAC sequencing using pooled methods. Methods Mol. Biol. 1227, 55–67 (2015).

91. Koren, S. et al. Canu: scalable and accurate long-read assembly via adaptive k-mer weighting and repeat separation. Genome Res. 27, 722–736 (2017).

92. Gordon, D., Abajian, C. & Green, P. Consed: a graphical tool for sequence finishing. Genome Res. 8, 195–202 (1998).

93. Campbell, M. S. et al. MAKER-P: a tool kit for the rapid creation, management, and quality control of plant genome annotations. Plant Physiol. 164, 513–524 (2014).

94. Cantarel, B. L. et al. MAKER: an easy-to-use annotation pipeline designed for emerging model organism genomes. Genome Res. 18, 188–196 (2008).

95. Kent, W. J. BLAT--the BLAST-like alignment tool. Genome Res. 12, 656–664 (2002).

96. Au, K. F., Underwood, J. G., Lee, L. & Wong, W. H. Improving PacBio long read accuracy by short read alignment. PLoS One 7, e46679 (2012).

97. Chaisson, M. J. & Tesler, G. Mapping single molecule sequencing reads using basic local alignment with successive refinement (BLASR): application and theory. BMC Bioinformatics 13, 238 (2012).

98. Kim, D. et al. TopHat2: accurate alignment of transcriptomes in the presence of insertions, deletions and gene fusions. Genome Biol. 14, R36 (2013).

99. The UniProt Consortium. UniProt: the universal protein knowledgebase. Nucleic Acids Res. 45, D158–D169 (2017).

100. O’Leary, N. A. et al. Reference sequence (RefSeq) database at NCBI: current status, taxonomic expansion, and functional annotation. Nucleic Acids Res. 44, D733–45 (2016).

101. Huerta-Cepas, J. et al. eggNOG 4.5: a hierarchical orthology framework with improved functional annotations for eukaryotic, prokaryotic and viral sequences. Nucleic Acids Res. 44, D286–93 (2016).

102. Huerta-Cepas, J. et al. Fast Genome-Wide Functional Annotation through Orthology Assignment by eggNOG-Mapper. Mol. Biol. Evol. 34, 2115–2122 (2017).

